# Induced sensitivity of *Bacillus subtilis* colony morphology to mechanical media compression

**DOI:** 10.1101/007773

**Authors:** Jessica K. Polka, Pamela A. Silver

## Abstract

Bacteria from several taxa, including *Kurthia zopfii*, *Myxococcus xanthus*, and *Bacillus mycoides*, have been reported to align growth of their colonies to small features on the surface of solid media, including anisotropies created by compression. While the function of this phenomenon is unclear, it may help organisms navigate on solid phases, such as soil. The origin of this behavior is also unknown: it may be biological (that is, dependent on components that sense the environment and regulate growth accordingly) or merely physical.

Here we show that *B. subtilis*, an organism that typically does not respond to media compression, can be induced to do so with two simple and synergistic perturbations: a mutation that maintains cells in the swarming (chained) state, and the addition of EDTA to the growth media, which further increases chain length. EDTA apparently increases chain length by inducing defects in cell separation, as the treatment has only marginal effects on the length of individual cells.

These results lead us to three conclusions. First, the wealth of genetic tools available to *B. subtilis* will provide a new, tractable chassis for engineering compression sensitive organisms. Second, the sensitivity of colony morphology to media compression in *Bacillus* is a physical rather than biological phenomenon dependent on a simple physical property of rod-shaped cells. And third, colony morphology under compression holds promise as a rapid, simple, and low-cost way to screen for changes in the length of rod-shaped cells or chains thereof.

## INTRODUCTION

Response of bacterial colony morphology (ie, orientation of growth) to small mechanical perturbations of growth media was first noted in *Kurthia*, a gram-positive genus notable for its striking feather-like morphology on gelatin slant cultures.(Sergent, 1906, 1907; Jacobsen, 1907; Stackebrandt, Keddie & Jones, 2006) A similar compression response has been reported in *Myxococcus xanthus*, where the phenomenon is dependent on adventurous motility, a flagellum- and pili-independent movement system.(Stanier, 1942; Fontes & Kaiser, 1999; Nan et al., 2014) Recently, the soil bacterium *Bacillus mycoides* was also shown to be sensitive to media perturbations.(Stratford, Woodley & Park, 2013) Interestingly, this compression response seems to occur by two different mechanisms: whereas individual *Myxococcus xanthus* dynamically reorients individual cells along lines of compression,(Dworkin, 1983) *Bacillus mycoides* instead gradually reorients the tips of chained cells as it grows.(Stratford et al., 2013)

The function of compression response is not known, but it has been suggested to aid navigation in natural environments on solid phases, like soil.(Dworkin, 1983) It has also been proposed as a potential tool for engineering applications in sensing environmental forces or generating patterns for nanofabrication.(Stratford et al., 2013)

Here we investigate whether increasing the length of chains of cells can induce compression sensitivity in an otherwise compression-insensitive species, *B. subtilis*. We employ a mutant of *B. subtilis* that forms long chains of cells (much like *B. mycoides*) and also deplete divalent cations in the media with EDTA; Mg^2+^ is thought be important for cell wall integrity. *B. subtilis* deprived of magnesium accumulates cell wall precursors,(Garrett, 1969) and magnesium is known to bind to components of the cell wall.(Heckels, Lambert & Baddiley, 1977) Notably, high magnesium concentrations can restore rod shape to cells with mutations in MreB, MreD, and PonA – all genes involved in cell wall synthesis.(Rogers, Thurman & Buxton, 1976; Rogers & Thurman, 1978; Murray, Popham & Setlow, 1998; Formstone & Errington, 2005)

## MATERIALS AND METHODS

### Time lapse microscopy

2% LB agar was cut into approximately 10mm × 10mm squares and inoculated with 1μl of liquid culture. The pad was then wedged, in a glass-bottomed dish (P35G-1.5-20-C, MatTek Corp.), between two plastic coverslips (Rinzl Plastic Coverslips, Size 22×22mm, Electron Microscopy Science) manually bent in half at a 90° angle. Thus, half of each plastic coverslip made contact with the bottom of the dish, while the other half made contact with the agar pad. After placing a drop of approximately 50μl of water on top of each plastic coverslip to maintain humidity in the dish, the MatTek dish was sealed with parafilm (this setup is illustrated in Fig. 1A). Cells were grown for approximately 6 hours at room temperature (approximately 23°) during a timelapse acquisition on a Nikon TE 2000 microscope equipped with an Orca ER camera, a 20x phase contrast objective, and Perfect Focus. A large area of the sample was composited with automatic image stitching by Nikon Elements AR. Areas toward the center of the pad were selected for imaging.

### Plate compression

Microtiter format plates were prepared with LB + 2% agar. 24 hours after plates were poured, sterilized polystyrene spacers (each 0.080” thick, for a total compression of 0.16” or 4.1 mm, equivalent to 4.8% compression) were inserted along the long dimension. Plates were stored at 37° for 24 hours, then inoculated from colonies grown on LB agar. Plates were incubated for 2-3 days at 30°, as the time required to reach colony dimensions >8 mm varied with EDTA concentration. After incubation, plates were imaged with a gel imager and colony dimensions measured with FIJI.(Schindelin et al., 2012)

### Cellular morphology

Colonies were grown on LB + 2% agar containing either 0 or 125μM EDTA. After 24 hours of incubation at 30°, cells from the edges of colonies were transferred directly to LB + 2% agar pads for imaging with the rounded bottoms of 0.6μl centrifuge tubes. To each pad, 1μl of an aqueous solution containing 10μg/ml FM4-64 (Invitrogen) was added. Cells were imaged with a 100X phase contrast objective, and cell and chain lengths were measured manually with spline-fitted segmented lines in FIJI. Two-sample KS tests were performed.(Kirkman, 1996)

## RESULTS

We first noted weak compression response of *B. subtilis* under the microscope. Unlike *B. mycoides*, *B. subtilis* colonies remain circular under compression under normal conditions. However, our microscopy assay (Fig. 1A) revealed that at small length scales (<100μm), *B. subtilis* cells display short-range alignment perpendicular to the direction of compression (marked with black arrows in Fig. 1A-C). Noting that the alignment is disrupted over longer length scales, we sought conditions under which *B. subtilis* cells might behave more similarly to *B. mycoides*. We noted that the chains of *B. subtilis* PY79 appeared shorter than that of *B. mycoides*, with the former reaching a maximum of approximately 300 µm (Fig. 1C), while the can extend for millimeters(Stratford et al., 2013).

**Figure 1.**
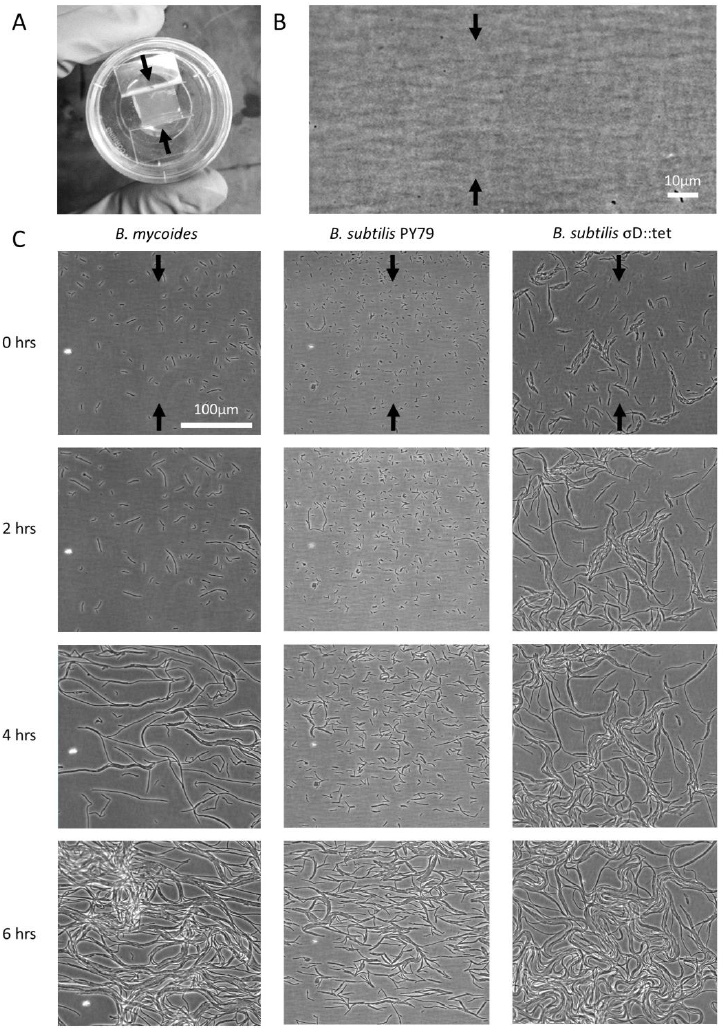
Microscopic morphology of *B. mycoides* and *B. subtilis* under compression. A) Cells from liquid culture were applied to the bottom of an agarose pad compressed between plastic coverslips in a MatTek dish. Black arrows indicate direction of compression throughout. B) Striations visible in agar surfaces. C) Montages of timelapses of *B. mycoides*, *B. subtilis* PY79, and *B. subtilis* σ^D^::tet. Note the striations visible in the agarose running perpendicular to the direction of compression.

To increase chain length, we used *B. subtilis* σ^D^::tet, a mutant that does not switch from swimming to swarming motility, and thus grows in long chains of cells (Kearns & Losick, 2005). To further perturb cell separation, we added EDTA to the growth medium.

To study colony morphology of *B. subtilis* under compression at the macroscopic scale with reproducible compression conditions, we prepared microtiter plates with LB + 2% agar and wedged polystyrene spacers between the agar and an edge of the plates (Fig. 2A). We inoculated the agar with colonies of *B. mycoides*, *B. subtilis* PY79, and *B. subtilis* σ^D^::tet. Under 4.8% compression, *B. mycoides* forms elongated colonies as reported,(Stratford et al., 2013) while, without EDTA, *B. subtilis* colonies are round (Fig. 2A). With the addition of EDTA to the media, both *B. subtilis* PY79 and σ^D^::tet display a compression response (Fig. 2B). This is dependent on the degree of compression; at 2.4% compression, both *B. subtilis* strains formed round colonies (data not shown).

**Figure 2.**
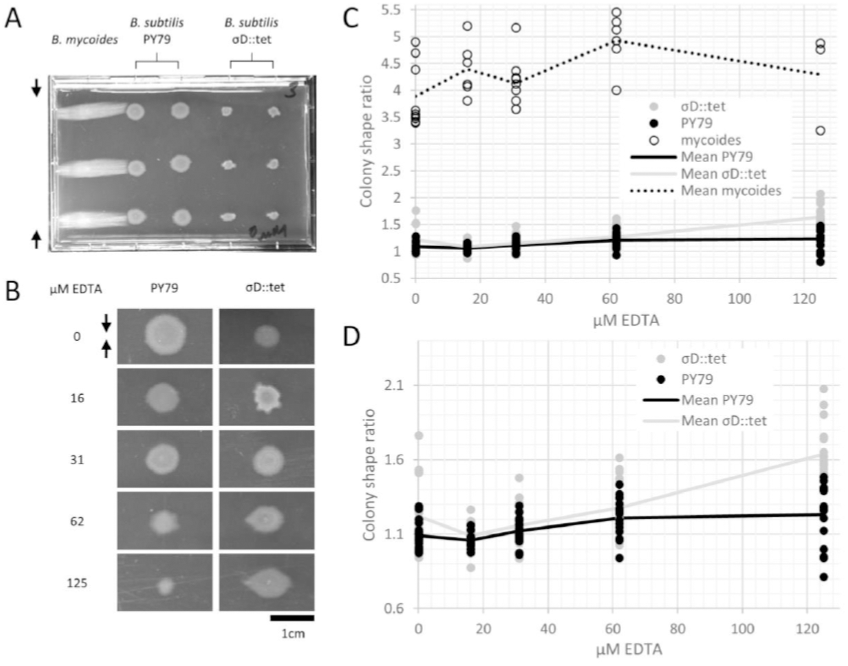
*B. mycoides* and *B. subtilis* colony morphology under compression. A) A microtiter plate inoculated with *B. mycoides* and *B. subtilis*. The two white bars at the top of the image of the plate are polystyrene spacers, totaling 4.8% of the plate height. Black arrows indicate direction of compression throughout. B) Representative images of *B. subtilis* PY79 and σD::tet colonies grown on compressed agar with varying EDTA concentrations. Scale bar, 1 cm. C) Plot of colony shape ratio (ie, colony measurement perpendicular to the dimension of compression/colony measurement parallel to the dimension of compression) as it varies with EDTA concentration. D) Same as in C but with axes scaled to emphasize relative effect of PY79 and σD::tet.

We next quantified this effect over several colonies under each EDTA condition at 4.8% compression. *Bacillus mycoides* forms colonies 4-4.5x larger in the dimension perpendicular to the direction of compression than parallel to it regardless of EDTA concentration (Fig. 2C). In comparison, the effect in *B. subtilis* is relatively small. *Bacillus subtilis* colonies were a maximum of approximately 1.5x larger in the direction perpendicular to compression, and this effect scaled with EDTA concentration (Fig. 2C). The EDTA effect was stronger for the σ^D^::tet strain; at 125uM EDTA, compressed σ^D^::tet colonies were 1.64x larger in the direction of compression (n = 17, standard deviation 0.21), while PY79 colonies were 1.23x larger (n = 16, standard deviation 0.20).

To understand how EDTA could affect compression response, we imaged cells taken directly from the edges of colonies on solid media containing either 0μM (Fig. 3A-C) or 125μM EDTA (Fig. 3D-F). The chains of *B. subtilis* cells, both PY79 and σ^D^::tet, are longer on 125μM EDTA, but cell lengths, as delineated by the membrane dye FM4-64, are only marginally different. Quantification of ∼300 chain and cell lengths for each strain under each condition (Fig. 4) reveals that *B. subtilis* chain lengths increase dramatically with the presence of EDTA, while *B. mycoides* chain lengths decrease slightly, suggesting that the EDTA effect on cell separation is specific to *B. subtilis* (Table 1).

**Figure 3.**
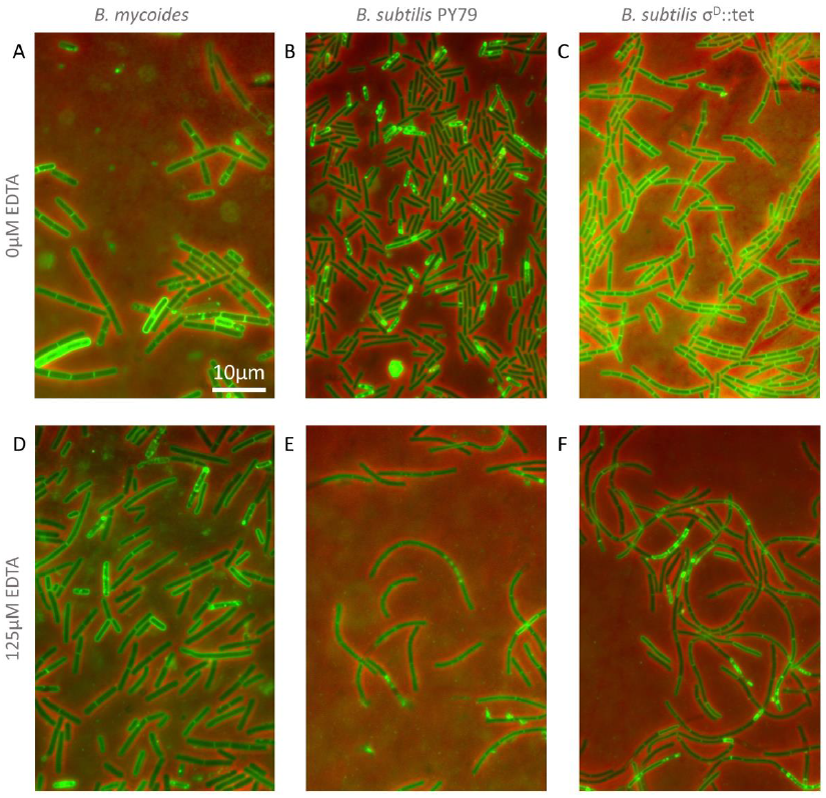
Cellular morphology with and without EDTA. A-C) *B. mycoides, B. subtilis* PY79, and *B. subtilis* σD::tet, respectively, growing on LB agar containing 0μM EDTA. D-F) As above on 125μM EDTA. In all images, phase contrast channel is in red, and FM4-64 is in green. Scale bar, 10μm.

**Figure 4.**
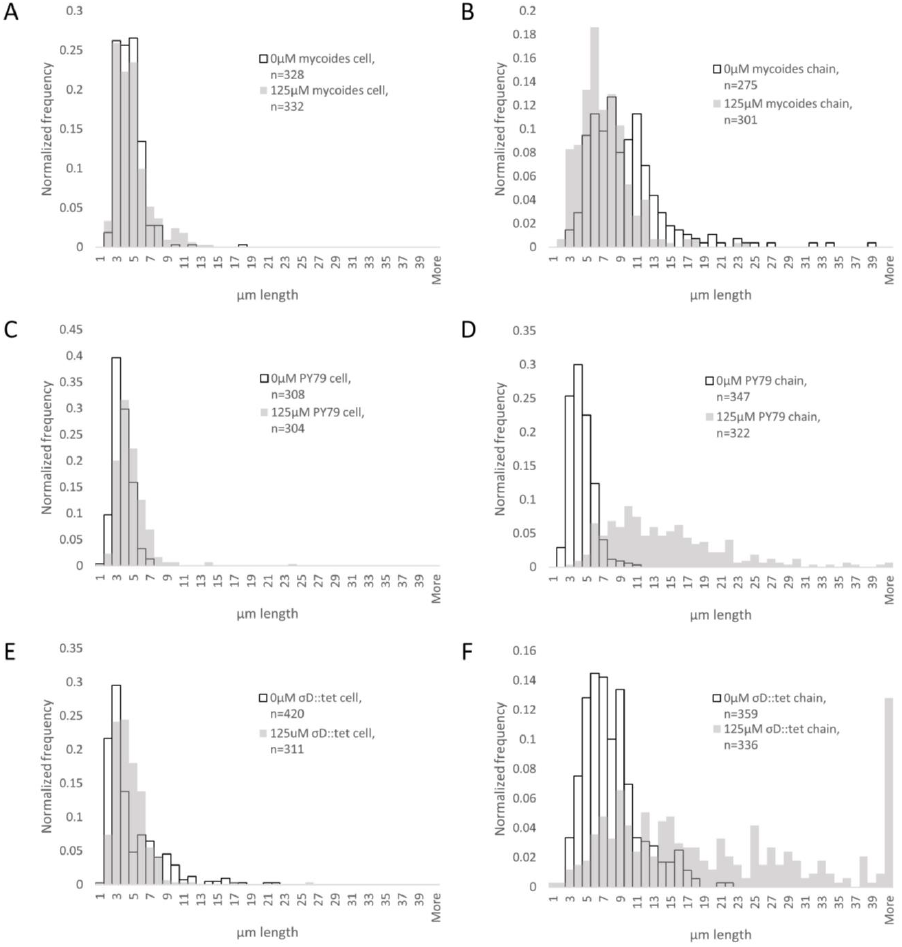
Quantification of chain and cell lengths with and without EDTA. A) Cell lengths of *B. mycoides* on 0μM (hollow bars) and 125μM EDTA (grey bars). B) Chain lengths of *B. mycoides*. C) Cell lengths of *B. subtilis* PY79. D) Chain lengths of *B. subtilis* PY79. E) Cell lengths of *B. subtilis* σD::tet. F) Chain lengths of *B. subtilis* σD::tet.

**Table 1.** Properties of cell and chain length measurement distributions

|  | Cell length |  |  | Chain length |  |  |
| --- | --- | --- | --- | --- | --- | --- |
|  | 0uM EDTA mean (μm) | 125uM EDTA mean (μm) | KS test maximum difference | 0uM EDTA mean (μm) | 125uM EDTA mean (μm) | KS test maximum difference |
| *B. mycoides* | 4.01 (st dev 1.54) | 4.33 (st dev 2.04) | D = 0.1044, P = 0.051 | 9.19 (st dev 4.81) | 6.60 (st dev 3.09) | D = 0.2959, P = 0.000 |
| *B. subtilis* PY79 | 3.18 (st dev 1.03) | 4.18 (st dev 1.93) | D = 0.2866, P = 0.000 | 3.94 (st dev 1.38) | 13.71 (st dev 7.23) | D = 0.8505, P = 0.000 |
| *B. subtilis* $\sigma^D::tet$ | 4.23 (st dev 3.20) | 4.12 (st dev 2.18) | D = 0.2413, P = 0.000 | 7.50 (st dev 3.36) | 21.99 (st dev 18.1) | D = 0.5633, P = 0.000 |

**Table 2.** Strains used in this study

| Designation | Description | Reference |
| --- | --- | --- |
| *B. subtilis* PY79 | Lab strain | Bacillus Genetic Stock Center 1A747 |
| *B. subtilis* $\sigma^D::tet$ | RL4169, DS323 | Kearns and Losick, 2005 (Kearns & Losick, 2005) |
| *B. mycoides* |  | ATCC 6462 |

## DISCUSSION

These results suggest that the phenomenon of colony orientation under compression can be induced in the model organism *B. subtilis*. In contrast to *Bacillus mycoides*, the genetic tractability of *B. subtilis* will facilitate engineering of compression sensitive bacteria for use as environmental sensors or guides for nanofabrication.(Stratford et al., 2013)

Furthermore, the fact that that colony orientation on compressed media is generalizable indicates that it is likely to be a physical phenomenon. Rather than requiring biological components specific to *B. mycoides*, it is probably based on factors like rod length, stiffness, and tip vs. isotropic growth pattern.

Long rod length is a common feature of two prototypical compression responders, *Bacillus mycoides* and *Kurthia sp.*, which both grow as long chains of cells.(Di Franco et al., 2002; Stackebrandt et al., 2006) As seen in microscopy of *B. mycoides*, the absence of cell separation allows the bacteria to find and maintain a direction of compression. This same chaining property is responsible for the baroque colony morphology of *B. mycoides*: mutants that do not display this colony morphology have shorter chain lengths.(Di Franco et al., 2002) Thus, compression response may be driven by the same mechanisms that influence colony morphology under normal conditions; these mechanisms influence the manner in which cells explore and colonize their environment, and may be of critical importance in soil environments.

In the case of *B. subtilis*, the increase in compression sensitivity is based on chain length (as a σ^D^ mutant responds more than PY79, and both respond more strongly in the presence of EDTA, which also increases rod length). Though EDTA likely affects multiple cellular processes, the role of Mg^2+^ in cell wall formation is clear.(Formstone & Errington, 2005) In particular, peptidoglycan hydrolases called autolysins are implicated in separation of cells after septation. Some of these autolysins, such as LytC, D, and F, are under the control of σ^D^.(Chen et al., 2009) However, LytC expression can also be driven by σ^A^,(Lazarevic et al., 1992) and this 50 kDa amidase is activated by addition of Mg2+ *in vitro*.(Foster, 1992) This magnesium dependence of LytC and its regulation by a second sigma factor may explain why EDTA treatment further increases chain length in σ^D^::tet cells. In addition to LytC, EDTA may be acting on other autolysins not regulated by σ^D^ (such as LytE or YwbG).(Smith, Blackman & Foster, 2000) The insensitivity of *B. mycoides* chain length to EDTA (Fig. 4, table 1) may be explained by species-specific differences in autolysins.

Inhibition of cell separation may not be the only relevant effect of EDTA, however. For example, perhaps depletion of Mg^2+^ changes the rigidity of cells such that they more readily align with the isotropic agar surface (Fig. 1B). An exhaustive understanding of EDTA’s effects on the mechanical properties of *B. subtilis* walls remains to be attained.

The relatively weak maximal compression response we achieved with *B. subtilis* compared to *B. mycoides* suggests that other factors limit the compression response of *B. subtilis*. We suggest that one contributing factor is the growth pattern of this organism. Whereas *B. mycoides* elongates from its tips,(Turchi et al., 2012) *B. subtilis* inserts cell wall isotropically along its length.(Tiyanont et al., 2006) In micrographs of *B. subtilis* under compression, the chains of cells appear more buckled than those of *B. mycoides* (Fig. 1C); perhaps friction prevents the distal ends of the chain from sliding along to accommodate new growth from the middle of the chain. This buckling disrupts adjacent chains and is likely to lead to a more disorganized colony morphology. In the future, further modifications, perhaps increasing surfactin production, may increase the magnitude of this response.

Finally, because *B. subtilis* compression response depends on chain length, we propose that under some circumstances, colony morphology under compression could serve as a simple, high-throughput assay for perturbations to bacterial cell length and chain formation.

## ACKNOWLEDGEMENTS

We thank Ethan Garner (Harvard University), Michael Baym (Harvard Medical School) and Ariel Amir (Harvard University) for helpful discussions. We are grateful to Stephanie Hays (Harvard Medical School) for critical reading of the manuscript.

## REFERENCES

Chen R, Guttenplan SB, Blair KM, Kearns DB. 2009. Role of the σD-Dependent Autolysins in Bacillus subtilis Population Heterogeneity. Journal of Bacteriology 191:5775–5784.

Dworkin M. 1983. Tactic behavior of Myxococcus xanthus. Journal of Bacteriology 154:452–459.

Fontes M, Kaiser D. 1999. Myxococcus cells respond to elastic forces in their substrate. Proceedings of the National Academy of Sciences 96:8052–8057.

Formstone A, Errington J. 2005. A magnesium-dependent mreB null mutant: implications for the role of mreB in Bacillus subtilis. Molecular Microbiology 55:1646–1657.

Foster SJ. 1992. Analysis of the autolysins of Bacillus subtilis 168 during vegetative growth and differentiation by using renaturing polyacrylamide gel electrophoresis. Journal of Bacteriology 174:464–470.

Di Franco C, Beccari E, Santini T, Pisaneschi G, Tecce G. 2002. Colony shape as a genetic trait in the pattern-forming Bacillus mycoides. BMC Microbiology 2:33.

Garrett AJ. 1969. The effect of magnesium ion deprivation on the synthesis of mucopeptide and its precursors in Bacillus subtilis. The Biochemical Journal 115:419–430.

Heckels JE, Lambert PA, Baddiley J. 1977. Binding of magnesium ions to cell walls of Bacillus subtilis W23 containing teichoic acid or teichuronic acid. Biochemical Journal 162:359–365.

Jacobsen H. 1907. Ueber einen richtenden Einfluss beim Wachstum gewisser Bakterien in Gelatine. Zentr. Bakt. Parasitenk. II:53–64.

Kearns DB, Losick R. 2005. Cell population heterogeneity during growth of Bacillus subtilis. Genes & Development 19:3083–3094.

Kirkman T. 1996. Statistics to use.

Lazarevic V, Margot P, Soldo B, Karamata D. 1992. Sequencing and analysis of the Bacillus subtilis lytRABC divergon: A regulatory unit encompassing the structural genes of the N-acetylmuramoyl-L-alanine amidase and its modifier. Journal of General Microbiology 138:1949–1961.

Murray T, Popham DL, Setlow P. 1998. Bacillus subtilis cells lacking penicillin-binding protein 1 require increased levels of divalent cations for growth. Journal of Bacteriology 180:4555–4563.

Nan B, McBride MJ, Chen J, Zusman DR, Oster G. 2014. Bacteria that Glide with Helical Tracks. Current Biology 24:R169–R173.

Rogers HJ, Thurman PF. 1978. Temperature-sensitive nature of the rodB maturation in Bacillus subtilis. Journal of Bacteriology 133:298–305.

Rogers HJ, Thurman PF, Buxton RS. 1976. Magnesium and anion requirements of rodB mutants of Bacillus subtilis. Journal of Bacteriology 125:556–564.

Schindelin J, Arganda-Carreras I, Frise E, Kaynig V, Longair M, Pietzsch T, Preibisch S, Rueden C, Saalfeld S, Schmid B et al. 2012. Fiji: an open-source platform for biological-image analysis. Nature Methods 9:676–682.

Sergent E. 1906. Des tropismes du “Bacterium zopfii” Kurth. Premiere note. Ann. inst. Pasteur:1005–1017.

Sergent E. 1907. Des tropismes du “Bacterium zopfii” Kurth. Deuxieme note. Ann. inst. Pasteur:842–850.

Smith TJ, Blackman SA, Foster SJ. 2000. Autolysins of Bacillus subtilis: multiple enzymes with multiple functions. Microbiology 146:249–262.

Stackebrandt E, Keddie RM, Jones D. 2006. The Genus Kurthia. In: Dr MDP, Falkow S, Rosenberg E, Schleifer K-H, Stackebrandt E eds. The Prokaryotes. Springer US, 519–529.

Stanier RY. 1942. A Note on Elasticotaxis in Myxobacteria. Journal of Bacteriology 44:405–412.

Stratford JP, Woodley MA, Park S. 2013. Variation in the Morphology of Bacillus mycoides Due to Applied Force and Substrate Structure. PLoS ONE 8:e81549.

Tiyanont K, Doan T, Lazarus MB, Fang X, Rudner DZ, Walker S. 2006. Imaging peptidoglycan biosynthesis in Bacillus subtilis with fluorescent antibiotics. Proceedings of the National Academy of Sciences of the United States of America 103:11033–11038.

Turchi L, Santini T, Beccari E, Di Franco C. 2012. Localization of new peptidoglycan at poles in Bacillus mycoides, a member of the Bacillus cereus group. Archives of Microbiology 194:887–892.

